# Pharmacological Modulation of TRIB2 Stability in Melanoma Reveals CDK12/13 as Dominant Regulators and Potential Therapeutic Targets

**DOI:** 10.1101/2024.07.17.603886

**Authors:** Victor Mayoral-Varo, Alba Orea-Soufi, Lucía Jiménez, Bruno Santos, Carlos Amenabar, Panagiota Kyriakou, Gisela Serrão, Miguel Jociles, Sara Obeso, Cristina Garnés-García, Pablo Fernández-Aroca, Ricardo Sánchez-Prieto, Bibiana I. Ferreira, Wolfgang Link

## Abstract

Tribbles homolog 2 (TRIB2) is one of three members of the Tribbles family of pseudo serine/threonine kinases. It acts as an oncogene whose expression is correlated with tumor progression and therapy resistance in melanoma. TRIB2 has been implicated in conferring resistance to various anti-cancer therapies suggesting TRIB2 as a therapeutic target for resistant tumors. This study explores the pharmacological targeting of TRIB2, revealing several independent routes of pharmacological manipulations. TRIB2 is a short-lived protein stabilized by inhibition of the PI3K/AKT pathway. Conversely, inhibitors of BRAF, MEK and ERK significantly decrease TRIB2 expression by a mechanism that involves transcription. Additionally, Polo-Like Kinases (PLKs) inhibitors significantly reduce TRIB2 protein expression and stability. Strikingly, increasing concentrations of the kinase inhibitor PIK-75 effectively eliminate TRIB2 in melanoma cells surpassing its PI3K inhibitory activity suggesting the destabilizing effect of the inhibition of an unknown kinase. TMT-based proteomics, chemical biology and genetic manipulation revealed CDK12 as the kinase that mediates the destabilizing effect on TRIB2. Overall, we identify three distinct classes of compounds that efficiently eliminate the oncogenic TRIB2 protein from melanoma cells based on different molecular mechanisms and exhibiting 40 to 200 times greater potency than the previously reported afatinib.

## Introduction

Tribbles homolog 2 (TRIB2) is the most ancestral member of the tribbles protein family, from which the other mammalian homologues TRIB1 and TRIB3 have evolved ^1^. Similar to other tribbles family members, TRIB2 possesses a conserved C-terminal motif that interacts with the ubiquitin E3 ligase machinery, along with a unique pseudocatalytic domain that lacks canonical metal-binding amino acids ^2^. Although the kinase activity of TRIB2 remains controversial, its role in promoting target protein degradation and regulating various signal transduction pathways has been well-established ^3^. Consequently, cells exhibit high sensitivity to changes in TRIB2 expression levels. Mechanistically, TRIB2 acts as an adaptor protein for substrate ubiquitination combining a kinase-like domain with a domain that facilitates interaction with ubiquitin ligases. While the pseudokinase domain is involved in substrate binding, the C-terminus of these proteins recruits the E3 ubiquitin-protein ligase COP1 leading to proteasomal degradation of the bZIP transcription factor C/EBPa ^4^. In melanoma, we have identified TRIB2 as a repressor protein of the FOXO3 transcription factor ^5^. The oncogenic role of TRIB2 stems from its function as a versatile scaffold protein that mediates the degradation of tumor suppressor proteins and the activation of tumor drivers. Overexpression of TRIB2 has been associated with malignant and/or chemoresistant phenotypes in various tumors, including acute myeloid leukaemia ^6^, chronic myelogenous leukemia ^7^, melanoma ^5,8,9^, small cell lung cancer ^10^, glioblastoma ^11^, colorectal cancer ^12,13^, and laryngeal squamous cell carcinoma ^14^. Recent findings also suggest that TRIB2 plays an important role in cancer immunotherapy, potentially protecting cold tumors from anti-cancer immunity ^15^. Taken together, the accumulating evidence supports TRIB2 as a therapeutic target for therapy-resistant tumors including melanoma ^16^. Additionally, TRIB2 has emerged as a druggable target ^17^ capable of binding ATP and undergoing autophosphorylation ^2^. Covalent inhibitors of the EGFR family such as the approved drug afatinib have been shown to bind to specific cysteine residues within the TRIB2 protein and induce its degradation ^18^. However, high concentrations of afatinib are required for effective TRIB2 degradation. Recently, Monga *et al* reported the direct binding of the antiviral drug daclatasvir to TRIB2 promoting its degradation through the proteasome ^13^. Daclatasvir was found to re-sensitizes enzalutamide-resistant prostate cancer cells to androgen receptor antagonist treatment. Moreover, a recent study identified nanobodies that bind the TRIB2 pseudokinase domain with low nanomolar affinity ^19^. The authors presented the first high-resolution structure of TRIB2 bound to a nanobody providing insights into its activated conformation. Advances in understanding TRIB2 regulation, structure and function lay the foundation for exploring various therapeutic strategies including structure-based drug design, ligand-based drug discovery as well as targeting TRIB2 expression and protein stability. In this study, we investigate TRIB2 stability and modulation of its expression using pharmacological approaches. Our findings demonstrate that TRIB2 protein is stabilized through the PI3K/AKT pathway activity, while its expression is downregulated by inhibition of components of the Mitogen11activated protein kinase (MAPK) pathway inhibition or silencing of Polo-Like Kinase 2 (PLK2) and CDK12/13.

## Results

### TRIB2 is rapidly degraded

To investigate the potential for targeting TRIB2 through pharmacological means, we initially conducted an analysis of TRIB2 stability and the mechanism of its turnover. Given that melanoma exhibits the highest expression of TRIB2 compared to other cancer types ^5^ (Figure S1A) and considering the significant role TRIB2 plays in the clinical outcome of melanoma, we specifically focused on measuring the half-life of TRIB2 in UACC-62 melanoma cells.

To block protein synthesis in these cells, we employed cycloheximide (CHX) treatment for 10 to 60 minutes. Following each 10-minute interval, we analysed the remaining amount of TRIB2 through western blotting. Figure 1A demonstrates that TRIB2 undergoes rapid degradation, with less than half of the protein remaining after 30 minutes of inhibiting new protein synthesis. Furthermore, after one hour of CHX treatment, almost no TRIB2 protein is detectable. Prolonged incubation with CHX for 2 to 8 hours renders TRIB2 undetectable (Figure S1B). Intriguingly, we observed that the inhibition of proteasomal degradation using the proteasome inhibitor MG132 led to increased TRIB2 protein suggesting that proteasomal degradation is involved in the process (Figure 1B).

**Figure 1.**
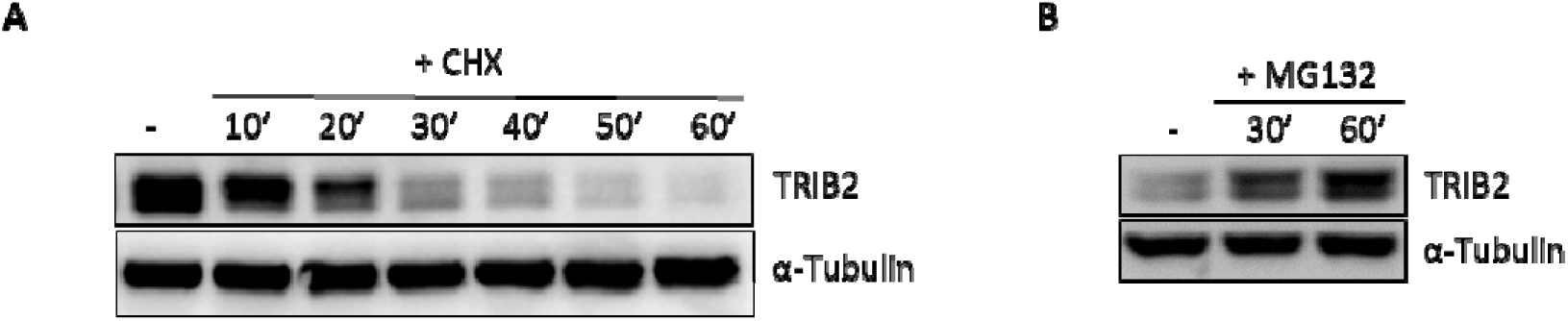
TRIB2 protein is rapidly degraded. **A)** Protein synthesis was blocked by Cycloheximide (CHX, 50 µg/mL) for the indicated time. The half-life of TRIB2 was measured by Western blot. **B)** Western blot analysis of TRIB2 protein levels following proteasome inhibition with 10 µM MG132 for the indicated time points. α-Tubulin was used as loading control.

### Induction of TRIB2 protein level upon PI3K/AKT pathway inhibition

To examine the impact of small molecule compounds that interfere with specific signal transduction pathways on TRIB2 protein amount, we used a collection of small molecule compounds with known mode of action. As afatinib was previously shown to bind to and promote the degradation of the TRIB2 protein in HeLa cells ^18^, we aimed to determine its effect in UACC-62 melanoma cells. The treatment of UACC-62 cells with afatinib require at least 10 µM for 4 hours to reduce TRIB2 protein level (Figure S2). These data indicate that covalent inhibitors of EGFR require optimization of their selectivity and potency for potential clinical exploitation of their anti-TRIB2 activity. Given our previous findings that TRIB2 influences PI3K/AKT signalling ^20^, we employed PI3K and mTOR inhibitors to treat UACC-62 cells and assessed their impact on the amount of TRIB2. As depicted in Figure 2, inhibition of the PI3K/AKT pathway led to an increase in TRIB2 protein levels. Treatment with the PI3K inhibitors PI-103 and LY294002, the dual PI3K/mTOR inhibitor dactolisib (BEZ-235), and the mTORC1 inhibitor rapamycin increased TRIB2 expression suggesting a reciprocal regulation between TRIB2 and AKT (Figure 2). Furthermore, we confirmed the inhibition of P70S6K phosphorylation in UACC-62 cells upon the exposure to these compounds (Figure 2). We also observed that the double band collapsed into a single band upon exposure to PI3K or mTOR inhibitors suggesting that direct phosphorylation of TRIB2 is involved in its destabilization. These data also could provide a mechanistic explanation of the paradoxical induction of TRIB2 upon the treatment with afatinib at 5 and 10 µM known to inhibit components of the PI3K/AKT signalling pathway.

**Figure 2.**
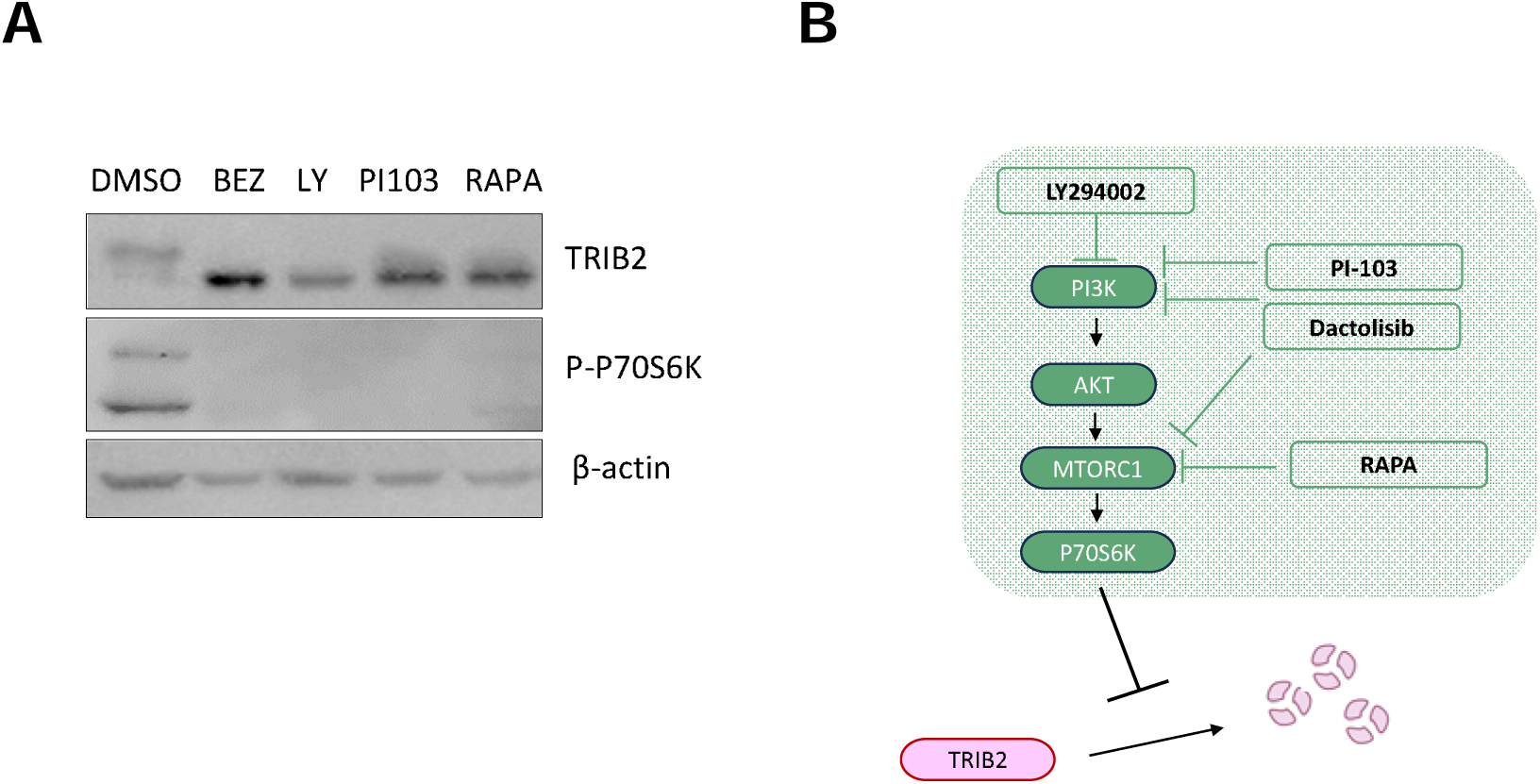
TRIB2 protein is regulated by the PI3K/AKT pathway. **A)** The treatment with dactolisib 500nM (BEZ), LY294002 25µM (LY), PI103 500nM or Rapamycin 5µM (RAPA) increases TRIB2 protein levels and decreases the level of P70S6K phosphorylation as determined by Western Blot. β-actin was used as loading control. **B)** Schematic model of the effect of different drugs including LY294002, PI-103, Dactolisib (BEZ), rapamycin (RAPA) on the PI3K pathway and TRIB2.

### MAPK pathway inhibition decreases TRIB2 expression

We also investigated the impact of inhibitors targeting the MAPK pathway, which plays a crucial role in melanoma and is frequently mutated in this type of cancer ^21^. Furthermore, TRIB2 has been shown to regulate MAPK signaling and bind MEK ^22^. Notably, treatment with vemurafenib, trametinib, and SCH-772984, inhibitors of BRAF, MEK and ERK respectively resulted in a decrease the level of endogenous TRIB2 protein (Figure 3A and Figure S2). To examine the kinetics of this effect, UACC-62 cells were treated with trametinib for 30 minutes, 1 hour, 2 hours, and 4 hours. No noticeable effect was observed at 30 minutes or 1 hour. However, levels began to decrease at 2 hours, and by 4 hours, TRIB2 protein was nearly undetectable (Figure 3B).

**Figure 3.**
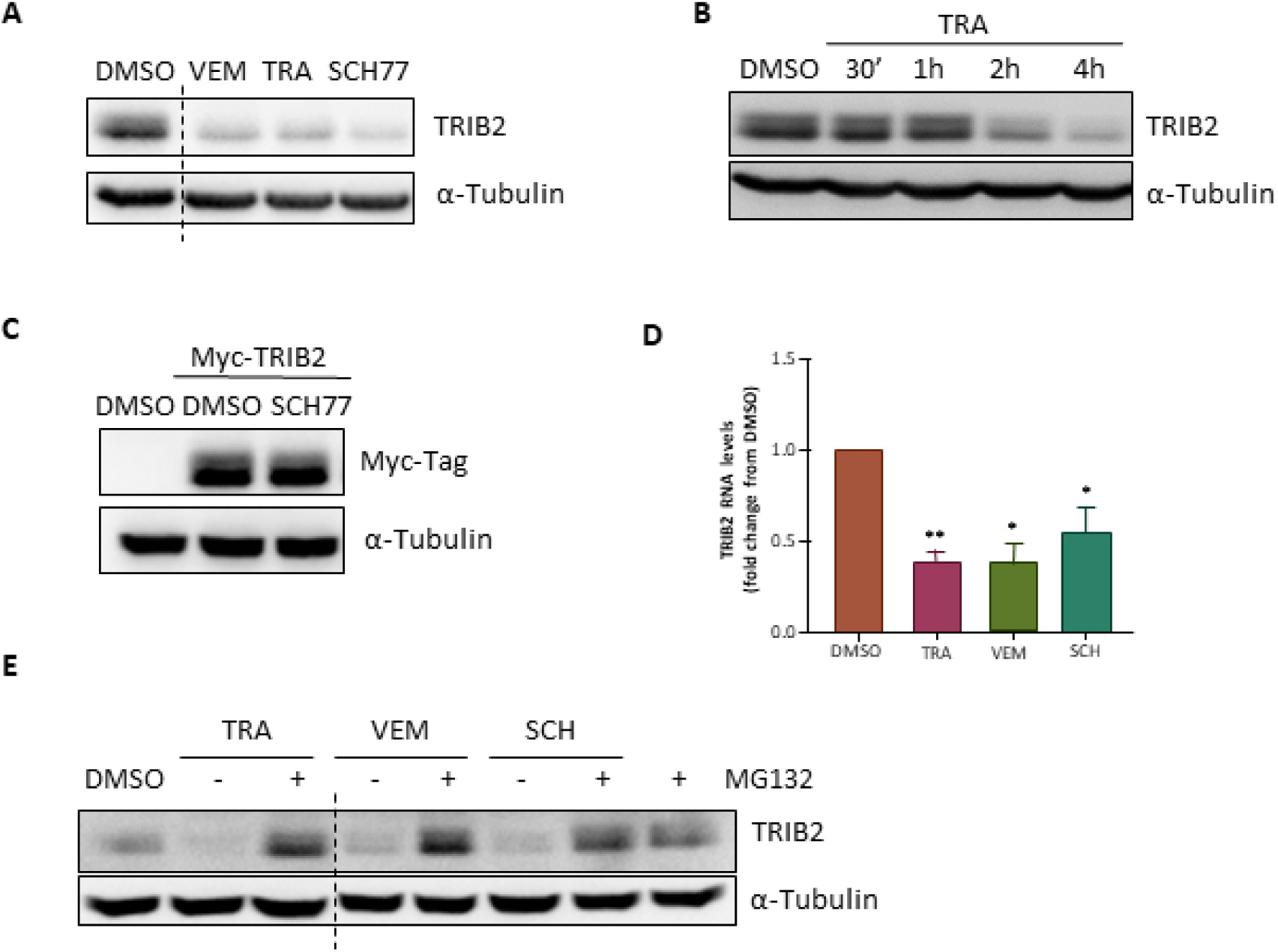
TRIB2 protein is downregulated by MAPK pathway inhibitors. **A)** Treatment with 500nM of vemurafenib (VEM), trametinib (TRA) or SCH-772984 (SCH77) reduces TRIB2 protein levels determined by Western Blot. **B)** Time-course of the effect of trametinib (500nM) treatment on TRIB2 protein levels at the indicated times measured by Western Blot. **C)** Transfected myc-TRIB2 is also downregulated by SCH-772984 (SCH77) treatment at 500nM 4 h **D)** Effect of Vemurafenib (VEM), Trametinib (TRA) or SCH-772984 (SCH) treatment for 4 h on TRIB2 RNA levels measured by qPCR. **E)** Treatment with the proteasomal inhibitor MG132 (10 µM) revert the downregulation of TRIB2 protein levels mediated by 500nM of tramentinib (TRA), vemurafenib (VEM) and SCH-772984 (SCH) treatment for 4 h. α-Tubulin has been used as loading control.

To investigate whether the inhibition of the MAPK pathway affectsTRIB2 at the protein level, we performed western blot analysis using cell lysates from cells ectopically expressing TRIB2 under the control of a constitutively active promoter. Figure 3C demonstrates that SCH-772984 did not affect the protein level of ectopically expressed TRIB2. Furthermore, we examined whether MAPK pathway inhibition could interfere with TRIB2 transcription. The results revealed that vemurafenib, trametinib and SCH-772984 downregulated the expression of TRIB2 transcripts (Figure 3D). In agreement, these MAPK inhibitors also decreased TRIB2 levels in A375 cells, indicating that the effect is not specific to UACC-62 cells (Figure S3). On the other hand, UACC-62 cells were co-treated with MAPK pathway inhibitors and the proteasome inhibitor MG132 for 4 hours. As shown in Figure 3E, in the presence of MG132 and MAPK pathway inhibitors, TRIB2 levels were higher than after DMSO vehicle or MG132 treatment alone. These findings suggest that inhibition of MAPK pathway components can act at both the RNA and the protein levels by promoting proteasome mediated degradation. In order to investigate whether the effect of MAPK pathway inhibitors on TRIB2 expression could be mediated through the regulation of downstream components of the pathway by ERK, we validated whether vemurafenib, trametinib and SCH-772984 affect the enzymatic activity of ERK. To this end we monitored the phosphorylation status RSK, a well-established substrate of ERK ^23^. Figure S2 demonstrates effective and potent inhibition of ERK and RSK phosphorylation upon exposure to the MAPK inhibitors. Additionally, we found that trametinib and vemurafenib failed to affect P70S6K phosphorylation and trametinib was unable to counteract P70S6K inhibition mediated by PI-103 (Figure S2). These data indicate that MAPK inhibitors effectively inhibit the enzymatic activity of ERK and act on TRIB2 levels independently of P70S6K. Conversely, we examined whether PIK-75 could affect MAPK signaling by monitoring the phosphorylation status of ERK and RSK. We showed that treatment of UACC-62 cells with PIK-75 failed to affect the phosphorylation of these proteins (Figure S2). Taken together, our results demonstrate that PIK-75 and MAPK inhibitors affect TRIB2 levels through two independent mechanisms.

### PLK inhibition abolishes TRIB2 expression destabilizing the protein

Intriguingly, the PLK1 inhibitor volasertib exhibited a time-dependent effect on TRIB2 expression. After just one hour of treatment, TRIB2 expression was significantly reduced, and it was eliminated after two hours (Figure 4A). We used a chemically similar derivative of volasertib, BI-2536 to test if we could confirm the effect of PLK1 inhibition on TRIB2 protein level. Similarly, to volasertib, BI-2536 significantly reduced TRIB2 protein level after 4 hours of treatment in a dose-dependent manner (Figure 4B). Notably, cells treated with 100nM BI-2536 exhibited significantly lower detectable levels of TRIB2 compared to cells treated with DMSO alone. Furthermore, TRIB2 levels continued to decrease at higher concentrations of BI-2536, namely 500nM and 1 µM. Similar to MAPK pathway inhibitors, volasertib did not modify the protein levels of TRIB2 whether its expression was driven by the endogenous or a constitutively active promoter (Figure 4C). These PLK inhibitors also led to a decrease in TRIB2 RNA levels (Figure 4D) suggesting that TRIB2 inhibition occurs at the transcriptional level. This mechanism of TRIB2 inhibition underscores the effectiveness of PLK inhibitors in regulating TRIB2 expression and highlights their potential as therapeutic agents for targeting TRIB2-mediated processes. To determine whether the effect of PLK inhibition on TRIB2 is dominant over PI3K inhibition we co-treated cells with volasertib or BI-2536 in combination with the potent panPI3K/mTOR inhibitor PI-103. As depicted in Figure 4E, the combined treatment also effectively reduced TRIB2 levels indicating that inhibition of PLKs completely overwrites the PI3K-mediated stabilization of TRIB2. To exclude the possibility that the effect of volasertib and BI-2536 is mediated through inhibition of the MAPK pathway, we measured the effect of these agents on the phosphorylation status of the ERK substrate RSK. As demonstrated in Figure S2, both inhibitors failed to affect RSK phosphorylation confirming that volasertib and BI2536 act independent of MAPK pathway inhibition.

**Figure 4.**
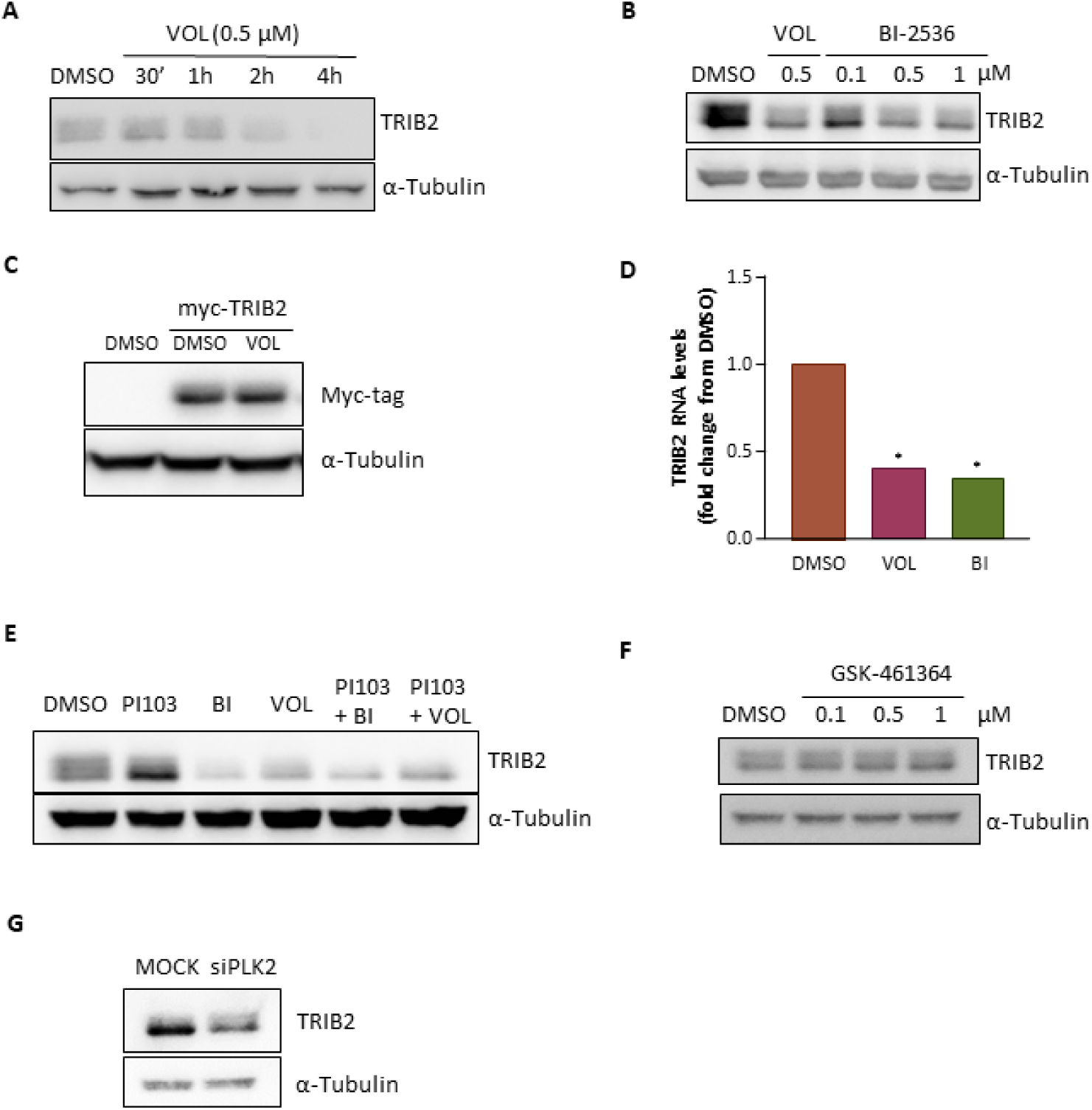
TRIB2 protein is downregulated by the PLK1 inhibitors Volasertib and BI-2536. A) Time-course of the effect of volasertib (VOL; 500nM) treatment on TRIB2 protein levels at the indicated times measured by Western Blot. B) Dose-response of BI-2536 treatment at 4h on TRIB2 protein levels measured by Western Blot. C) Transfected myc-TRIB2 is not downregulated by volasertib (VOL) at 500nM 4h. D) Effect of volasertib and BI-2536 treatment for 6h on TRIB2 RNA levels measured by qPCR. E) Effect of the combination of volasertib (VOL) and BI-2536 (BI) with the PI3K inhibitor (PI-103) on TRIB2 levels determined by Western Blot. F) Dose-response of GSK-461364 treatment at 4h on TRIB2 protein levels measured by Western Blot. I) Effect of PLK2 downregulation using esiRNA technology on TRIB2 protein levels determined by Western Blot.

As volasertib and BI-2536 potently inhibit not only PLK1 but also PLK2 and PLK3, albeit with slightly lower potency but still in the lower nanomolar range, we aimed to identify which PLK isoform is primarily responsible for the effect on TRIB2. We investigated the ATP-competitive PLK1 inhibitor GSK-461364, which exhibits a 390-fold greater selectivity for PLK1 compared to PLK2 and PLK3, and over 1,000-fold selectivity relative to a panel of 48 other kinases. Interestingly, GSK-461364 failed to affect TRIB2 levels (Figure 4F) suggesting that TRIB2 stability is regulated by PKL2 or 3 rather than PLK1. Additionally, wortmannin known to potently inhibit PLK1 and PLK3 ^24^ actually increased TRIB2 protein levels due to its PI3K inhibitory activity (data not shown). Collectively, these findings suggest that PLK2 is likely the key enzyme responsible for mediating the effect of volasertib and BI-2536 on TRIB2. To verify our hypothesis, we silenced PLK2 and measure its effect on TRIB2 protein levels. As shown in Figure 4G, transient PLK2 silencing reduced TRIB2 levels indicating that the inhibition of PLK2 promotes TRIB2 downregulation and thus, supports that the effect of volasertib and BI-2536 on TRIB2 is mediated by PLK2.

### PIK-75 eliminates TRIB2 acting at the protein level

Interestingly, unlike other PI3K inhibitors, increasing nanomolar concentrations of PIK-75 resulted in a reduction of TRIB2 protein levels after 4 hours of treatment. Notably, 200nM of PIK-75 was sufficient to override the stabilizing effect, leading to a significant decrease in the amount of TRIB2 protein. Treatment with 500nM PIK-75 virtually eliminates TRIB2 from UACC-62 cells (Figure 5A). The decline in TRIB2 expression began as early as one hour into PIK-75 treatment and reached complete elimination after 2 hours (Figure 5B). To investigate whether the decrease in TRIB2 protein was due to suppressed TRIB2 transcription, we performed qPCR analysis. Treatment with 500nM PIK-75 only resulted in a minor reduction of TRIB2 transcripts (Figure 5C). To further confirm that PIK-75-mediated elimination of TRIB2 in melanoma cells primarily occurred at the protein level, we ectopically expressed Myc-tagged TRIB2 transcribed from a constitutively active promoter. Figure 5D demonstrates that PIK-75 affected the stability of the ectopically expressed TRIB2 protein suggesting that its action is independent of transcription. In order to examine whether drug-induced TRIB2 destabilization is a cell-specific phenomenon, we treated different cell lines with small molecule compounds. In particular, we used A375 melanoma cells and osteosarcoma U2OS cells stably expressing TRIB2-GFP to PIK-75. As shown in Figure S3 endogenous TRIB2 in A375 cells decreased upon PIK-75 exposure. Similarly, PIK-75 also affected TRIB2 level when ectopically expressed from a constitutively active gene promoter. These data indicate that PIK-75 treatment exerts its effect independent of the cell type and affects TRIB2 independently of transcription at the protein level. Interestingly, co-treatment of UACC-62 cells with MG132, a proteasome inhibitor, and 500nM PIK-75 only modestly rescued TRIB2 levels compared to cells treated with DMSO vehicle control indicating that decrease in TRIB2 induced by PIK-75 is partly independent of proteasomal degradation (Figure 5E). To determine whether the negative regulation induced by 500nM PIK-75 dominates over the positive effect of PI3K inhibition, we combined PIK-75 treatment with the exposure to the potent panPI3K/mTOR inhibitor PI-103. As shown in Figure 5E, in the presence of 500nM PIK-75, PI-103 is unable to counteract the PIK-75-mediated decrease of TRIB2 protein. We hypothesized that as the concentration of PIK-75 increases, its specificity for its primary target PI3K, may diminish. As PIK-75 might directly bind to TRIB2 and destabilize it, we performed Cellular Thermal Shift Assays (CETSAs) which involve subjecting cell lysates to increasing temperatures and quantifying the presence of the protein of interest in the soluble fraction through immunoblotting. Interestingly, TRIB2 exhibited faster solubility loss upon heating compared to the control proteins PI3K and α-tubulin (Figure 5F). While PI3K displayed stabilization in the presence of PIK-75 at 55°C, no detectable difference was observed in TRIB2 with or without PIK-75. These findings suggest that PIK-75 does not directly bind to TRIB2.

**Figure 5.**
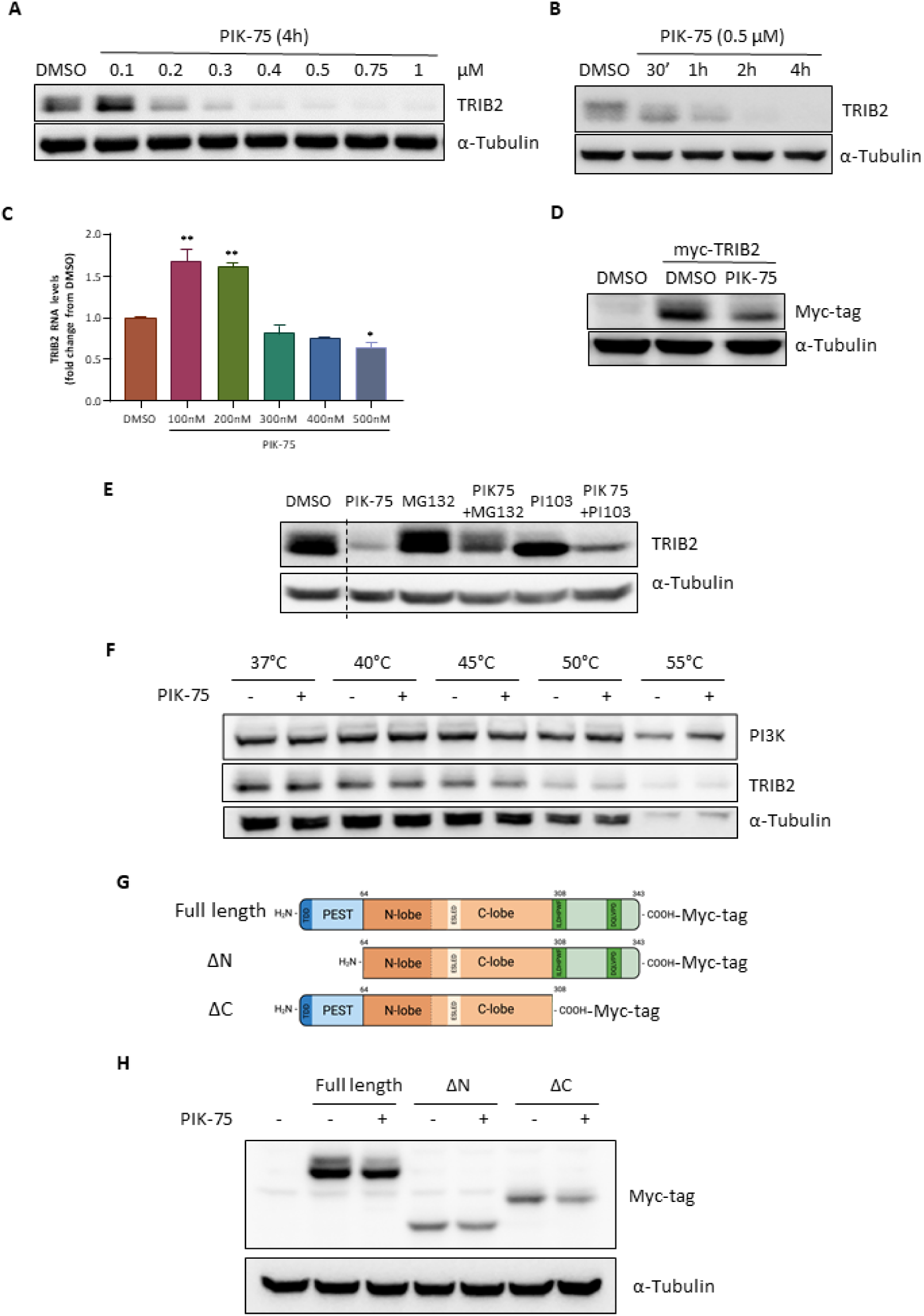
TRIB2 protein is downregulated by PIK-75. **A)** Time-course of the effect of PIK-75 (500nM) treatment on TRIB2 protein levels at the indicated times measured by Western Blot. **B)** Dose-response of PIK-75 treatment at 4h on TRIB2 protein levels measured by Western Blot. **C)** Effect of PIK-75 treatment at 4h on TRIB2 RNA levels measured by qPCR. **D)** Transfected myc-TRIB2 is also downregulated by PIK-75 treatment at 500nM and 4h. **E)** Effect of the combination of PIK-75 with the cell-permeable proteasome inhibitor MG132 or the PI3K inhibitor PI-103 on TRIB2 levels determined by Western Blot. **F)** Effect of 30 minutes PIK-75 (100 µM) treatment at the indicated temperatures on TRIB2 levels. PI3K has been used as a positive control of the stabilizing effect of PIK-75. **G)** Representation of the full length, and the deletion mutants ΔN and ΔC of TRIB2. **H)** Effect of PIK-75 on TRIB2 deletions mutants. α-Tubulin has been used as loading control.

### The N-terminus of TRIB2 is necessary for PIK-75-mediated TRIB2 destabilization

To identify the specific region of the TRIB2 protein responsible for the destabilizing effect of PIK-75, we employed deletion mutants of TRIB2 genetically fused to a Myc-tag as previously described ^25^. We transfected the full-length construct and the derived deletion mutants (Figure 5G) into UACC-62 cells. Subsequently, we analyzed the ratio of TRIB2 protein levels between vehicle-treated cells and cells exposed to 500nM PIK-75 for each deletion mutant using western blotting. Interestingly, only the N-terminal deletion protected TRIB2 from PIK-75-mediated degradation while the C terminal deletion remained susceptible to the effects of PIK-75 (Figure 5H). These results suggest that the N-terminus of TRIB2 contains critical amino acids that may undergo posttranslational modifications contributing to the stabilization of TRIB2 protein, a process that is abolished upon PIK-75 treatment.

### Isoform-specific effect of TRIB2 inhibitors on Tribbles proteins

We proceeded to examine the impact of the identified TRIB2 inactivating compounds on other members of the Tribbles protein family, TRIB1 and TRIB3, in melanoma cells. The cells were treated with the previously used inhibitors: PI-103 (PI3K/mTOR), SCH-772984 (ERK), PIK-75 (multi-kinase), BI-2536, GSK-461364 and volasertib (PLKs) for a duration of 4 hours. Figure S4 illustrates that none of the compounds produce a decrease in the quantity of TRIB1. However, in the case of TRIB3, PI-103, BI-2536 and volasertib downregulate its amount in the treated cells. Intriguingly, PIK-75 failed to exert an effect on TRIB1 or TRIB3 indicating its specificity towards TRIB2.

To determine whether the influence of ERK or PLK inhibition on TRIB3 levels occurs at the protein level, we treated cells that produced TRIB3 from a constitutive promoter. Figure S4 demonstrates that the quantity of ectopic TRIB3 remained unchanged following any of the treatments. This suggests that the agents affect TRIB3 through a mechanism that involves transcription, implying that the posttranslational regulation observed for TRIB2 is specific to this particular isoform.

### Phosphoproteomics identifies hypophosphorylated peptides and proteins

We consider the finding that PIK-75 can completely eliminate the TRIB2 oncoprotein from melanoma cells within one to four hours, at a concentration at least 100-fold more potent than the previously reported afatinib clinically highly relevant. Accordingly, we sought to characterize the mechanism by which PIK-75 exerts this effect on TRIB2. To identify the molecular target responsible for the PIK-75–induced destabilization of TRIB2, we conducted Tandem Mass Tag (TMT)-based quantitative total and phospho-proteomic analyses. The experimental workflow is depicted in Figure 6A. UACC-66 wt cells were exposed to 500nM of PIK-75 or vehicle (DMSO) for one hour. Four replicates per condition were used to extract proteins followed by precipitation, reduction, cysteine alkylation, and tryptic digestion. Isobaric labeling was performed using TMT reagents, labeling 50 µg of peptides per condition along with an internal standard (IS).

**Figure 6.**
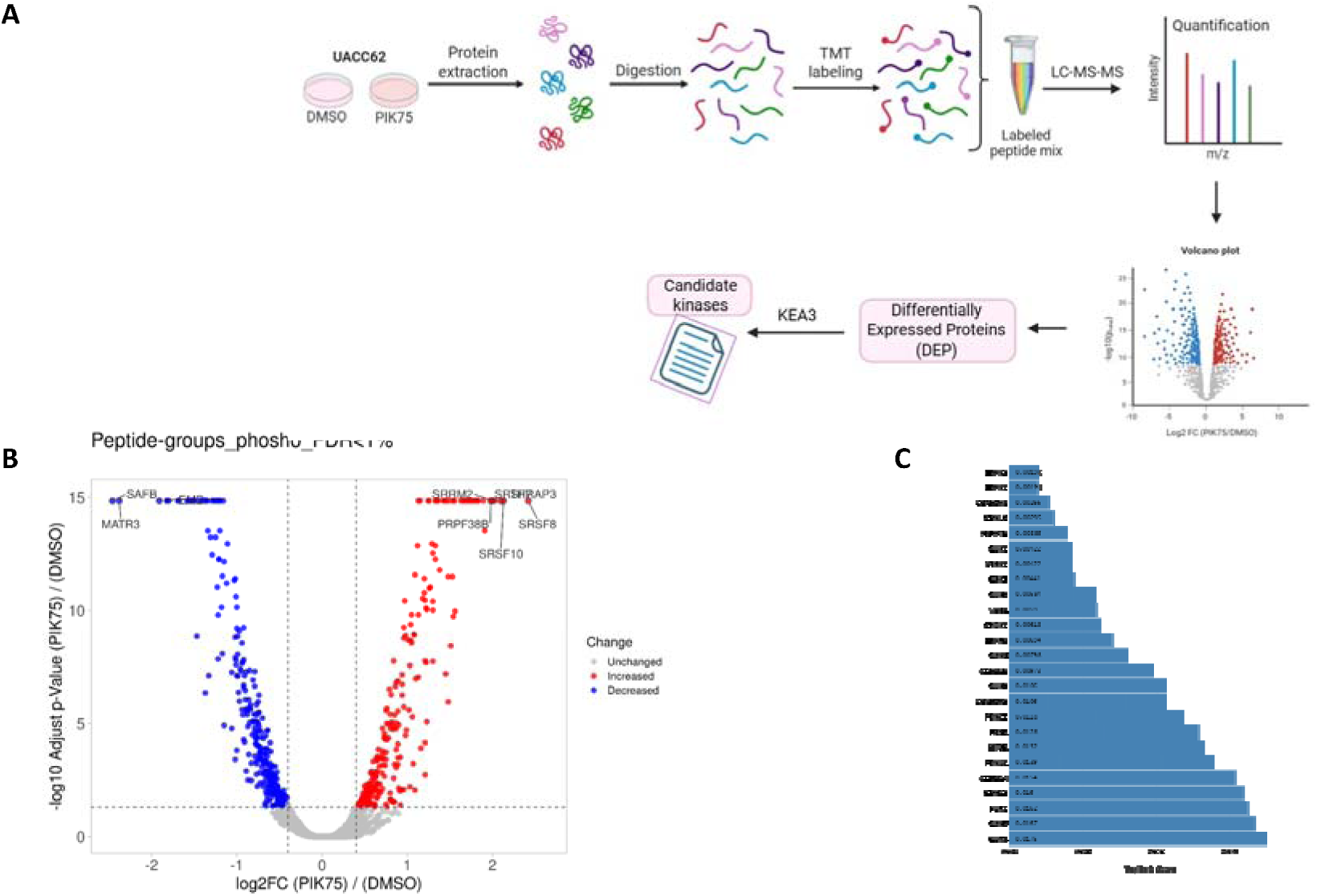
Phosphoproteomics analysis reveals candidate Kinases inhibited by PIK-75. **A)** Schematic workflow of the study Experimental parameters of the analysis: UACC62 w.t. DMSO/PIK-75 (500nM) – 4h. The illustrations to show the preparation steps of the analysis: Lysis, Protein precipitation, Protein Quantification, Reduction, alkylation of cysteines, and insolution tryptic digestion, Isobaric labeling (TMT reagent), Purification and enrichment of phosphopeptides, Purification, desalting, and removal of interferents. **B)** Volcano plot of the phosphopeptides according to the average ratio of three technical replicates and p value (−L*log10 p value). Grey points represent unchangedphosphopeptides with a log2 fold change (log2FC) value between −0.4 and 0.4 (changed <31%) and/or a p-value >0,05, red and blue represent the upregulated and downregulated proteins respectively, with a log2 fold change (log2FC) value <-0.4 or >0.4 and a p-value <0,05; the horizontal dashed line indicates p value*L*=*L*0.05. **C)*** TopRank bar chart displays the TopRank score of the highest-ranked substrates of the kinases in the analyzed list. Generated using the KEA3 app.

A total of 5289 unique proteins and 61 redundant protein candidates were identified in the total proteome (FDR <1%), while 3129 proteins and 87 candidates were identified in the phosphoproteomic analysis. Relative quantification using TMT labeling revealed 39 differentially expressed proteins (adjusted p-value <0.05) in the total proteome, with only 17 proteins showing robust differential expression when filtered for peptides >2 indicating that one-hour treatment with PIK-75 exposure does not significantly affect total protein expression, as expected. In the phosphoproteomic dataset, 7573 phosphopeptides were quantified, corresponding to 2339 unique phosphoproteins. A total of 582 phosphopeptides were differentially regulated (adjusted p-value <0.05), with phosphorylation site probabilities provided for high-confidence assignment (>75%) Figure S5A and Table S1).

Protein abundance ratios (PKI75/DMSO) were normalized to the internal standard. Log_2_ transformation was applied for symmetrical interpretation of up- and downregulated proteins. Statistical significance was determined using a t-test with Benjamini-Hochberg correction to control the false discovery rate. Principal component analysis (PCA) demonstrated clear clustering of biological replicates for both total and phosphoproteomic datasets, confirming data quality and reproducibility (Figure S5B). Volcano plots representing fold change distribution of phosphopeptides and phosphoproteins identified in UACC62 cells treated with DMSO or PIK-75 are depicted in Figure 6B and C respectively.

### Phosphoproteomic analysis reveals candidate kinases inhibited by PIK-75

To identify the kinases potentially responsible for the phosphorylation of hypophosphorylated peptides and proteins upon PIK-75 treatment, we performed Kinase Enrichment Analysis 3 (KEA3). KEA3 determines the overrepresentation of upstream kinases whose known substrates overlap with the list of differentially phosphorylated proteins (Figure 6C). Additionally, we used the kinase prediction feature to query the PhosphoSitePlus database for candidate kinases likely responsible for the observed hypophosphorylation events in PIK-75-treated UACC62 cells (Table S2). Notably, the top-ranked candidate kinases identified by both approaches were predominantly members of the CMGC family of serine/threonine protein kinases, including Glycogen Synthase Kinase 3 (GSK3α/β), CDC-like kinases (Clk1/Sty), Serine/Arginine Protein Kinases (SRPK1/2), and Cyclin-Dependent Kinases (CDKs).

**Table S2.** List of possible kinases generated from KEA3 app and Phosphosite plus: KEA3 Results with High-Moderate Score. Mean score L 120 = Weak.

| Rank | Protein | Integrated Scaled Rank | Overlapping Proteins |
| --- | --- | --- | --- |
| 1 | SRPK2* | 3.5 | 97 proteins |
| 2 | CLK1* | 5.714 | 73 proteins |
| 3 | SRPK3 | 6.4 | 38 proteins |
| 4 | CLK2 | 12.56 | 69 proteins |
| 5 | SRPK1* | 15.4 | 80 proteins |
| 6 | CDK9 | 30.91 | 61 proteins |
| 7 | CDK2* | 36 | 110 proteins |
| 8 | CSNK2A1* | 38.27 | 133 proteins |
| 9 | CDK1 | 39.8 | 115 proteins |
| 10 | CSNK2A2 | 42.8 | 69 proteins |
| 11 | ABL1 | 43.27 | 76 proteins |
| 12 | AURKB | 43.45 | 73 proteins |
| 13 | CLK3 | 44.67 | 52 proteins |
| 14 | ATM | 49.64 | 101 proteins |
| 15 | PLK1 | 49.64 | 80 proteins |
| 16 | SRC | 49.91 | 118 proteins |
| 17 | SCYL2 | 53 | 39 proteins |
| 18 | AKT1 | 54.36 | 129 proteins |
| 19 | CHEK2 | 56 | 66 proteins |
| 20 | CDK6 | 56.9 | 72 proteins |
| 21 | GSK3B | 58.73 | 114 proteins |
| 22 | MTOR | 59.27 | 91 proteins |
| 23 | AURKA | 64.09 | 63 proteins |
| 24 | DYRK1A | 66.64 | 49 proteins |
| 25 | MAPK1 | 68.45 | 105 proteins |
| 26 | PRKDC | 71.64 | 62 proteins |
| 27 | CSNK2A3 | 75 | 39 proteins |
| 28 | MAPK3 | 75.36 | 104 proteins |
| 29 | ROCK1 | 77 | 67 proteins |
| 30 | CLK4 | 79.83 | 47 proteins |
| 31 | PRKCA | 82.18 | 96 proteins |
| 32 | RPS6KB1 | 83 | 72 proteins |
| 34 | CSNK1E | 83.55 | 53 proteins |
| 35 | PRPF4B | 83.67 | 65 proteins |
| 36 | PRKACA | 84.27 | 102 proteins |
| 37 | EGFR | 84.55 | 95 proteins |
| 38 | PRKAA1 | 86.3 | 65 proteins |
| 39 | MKNK1 | 86.78 | 17 proteins |
| 40 | PAK1 | 87.27 | 65 proteins |
| 41 | PRKCI | 88.82 | 70 proteins |
| 42 | LATS2 | 89.11 | 26 proteins |
| 43 | RPS6KA1 | 89.9 | 48 proteins |
| 44 | CDK11A | 90 | 42 proteins |
| 45 | CDC7 | 90.44 | 25 proteins |
| 46 | CDK8 | 91.8 | 32 proteins |
| 47 | FYN | 92.55 | 78 proteins |
| 48 | RIOK2 | 94.83 | 30 proteins |
| 49 | MARK2 | 94.91 | 43 proteins |
| 50 | MAPK14 | 95.27 | 89 proteins |
| 51 | CSNK1A1 | 95.91 | 59 proteins |
| 52 | STK4 | 97.67 | 46 proteins |
| 53 | PRKCZ | 97.82 | 66 proteins |
| 54 | CHEK1 | 98 | 67 proteins |
| 55 | LRRK2 | 98.44 | 77 proteins |
| 56 | ATR | 98.6 | 65 proteins |
| 57 | MAP2K1 | 99.36 | 44 proteins |
| 58 | MAPK13 | 99.56 | 43 proteins |
| 59 | CDK4 | 101.2 | 86 proteins |
| 60 | GSK3A | 101.9 | 52 proteins |
| 61 | CDK7 | 102 | 59 proteins |
| 62 | TTN | 102.5 | 34 proteins |
| 63 | MAPK6 | 102.7 | 56 proteins |
| 64 | BUB1 | 103.3 | 41 proteins |
| 65 | PDPK1 | 103.4 | 66 proteins |
| 66 | LATS1 | 103.7 | 39 proteins |
| 67 | MYLK | 104.3 | 23 proteins |
| 68 | TESK2 | 105.4 | 31 proteins |
| 69 | ILK | 106.8 | 50 proteins |
| 70 | PDK2 | 107.8 | 11 proteins |
| 71 | PTK2 | 108 | 44 proteins |
| 72 | HIPK2 | 109.2 | 52 proteins |
| 73 | TNK2 | 109.5 | 33 proteins |
| 74 | CDK12 | 109.8 | 38 proteins |
| 75 | NTRK1 | 109.9 | 99 proteins |
| 76 | WNK1 | 110.1 | 17 proteins |
| 77 | ERN1 | 111 | 31 proteins |
| 78 | CSNK1D | 112.9 | 44 proteins |
| 79 | CSK | 113.9 | 41 proteins |
| 80 | JAK2 | 115.2 | 70 proteins |
| 81 | YES1 | 116.3 | 47 proteins |
| 82 | MAPK8 | 116.5 | 102 proteins |
| 83 | IKBKE | 116.7 | 28 proteins |
| 84 | EIF2AK2 | 117 | 36 proteins |
| 85 | RIOK1 | 117.9 | 36 proteins |
| 86 | RIPK3 | 118.8 | 21 proteins |
| 87 | PRKCB | 119.4 | 37 proteins |

### Chemical biological validation of the candidate kinases

To further evaluate the functional relevance of the predicted kinases, we investigated the effects of small-molecule inhibitors targeting members of the CMGC kinase family. Several well-characterized inhibitors are available including, TWS119, CHIR-98014 and Laduviglusib (targeting GSK3), TG003 (targeting Clk1), and SPHINX31 (targeting SRPK1/2). These compounds serve as valuable tools to assess whether inhibition of the predicted kinases can recapitulate or modulate the phospho-signaling changes induced by PIK-75. However, none of these inhibitors significantly affected TRIB2 levels (Figure S6).

In contrast, SR-4835—a potent inhibitor of CDK12/13—dramatically reduced TRIB2 protein levels (Figure 7). THZ531, another CDK12/13 inhibitor, also reduced TRIB2 expression (Figure S6). To determine whether this effect might be due to general CDK inhibition, we used Palbociclib, a specific inhibitor of CDK4/6. Palbociclib treatment did not alter TRIB2 levels (Figure S7A), suggesting that the effect is specific to CDK12/13 inhibition and may be the mechanism by which PIK-75 downregulates TRIB2. To investigate whether CDK12/13 inhibition recapitulates key aspects of PIK-75-mediated TRIB2 regulation, we conducted dose–response and time-course experiments using SR-4835. Similar to PIK-75, SR-4835 rapidly decreased TRIB2 levels (Figure S7B). We then treated UACC-62 cells with four different concentrations of SR-4835 for four hours and assessed TRIB2 protein levels using western blot analysis. As shown in Figure S7C, TRIB2 levels began to decrease at 2 µM, and treatment with 10 µM almost completely eliminated TRIB2 from the cells. We next asked whether SR-4835 can also reduce TRIB2 protein levels when TRIB2 is expressed from a constitutively active promoter, as previously shown for PIK-75. To test this, we used U2OS cells, which lack endogenous TRIB2 expression, and ectopically expressed GFP-tagged TRIB2 under the control of the human cytomegalovirus immediate-early promoter—a strong and constitutive promoter. Upon treatment with SR-4835 or PIK-75 for 4 hours, we observed a strong and comparable reduction in TRIB2 protein levels for both compounds (Figure S7D).

**Figure 7.**
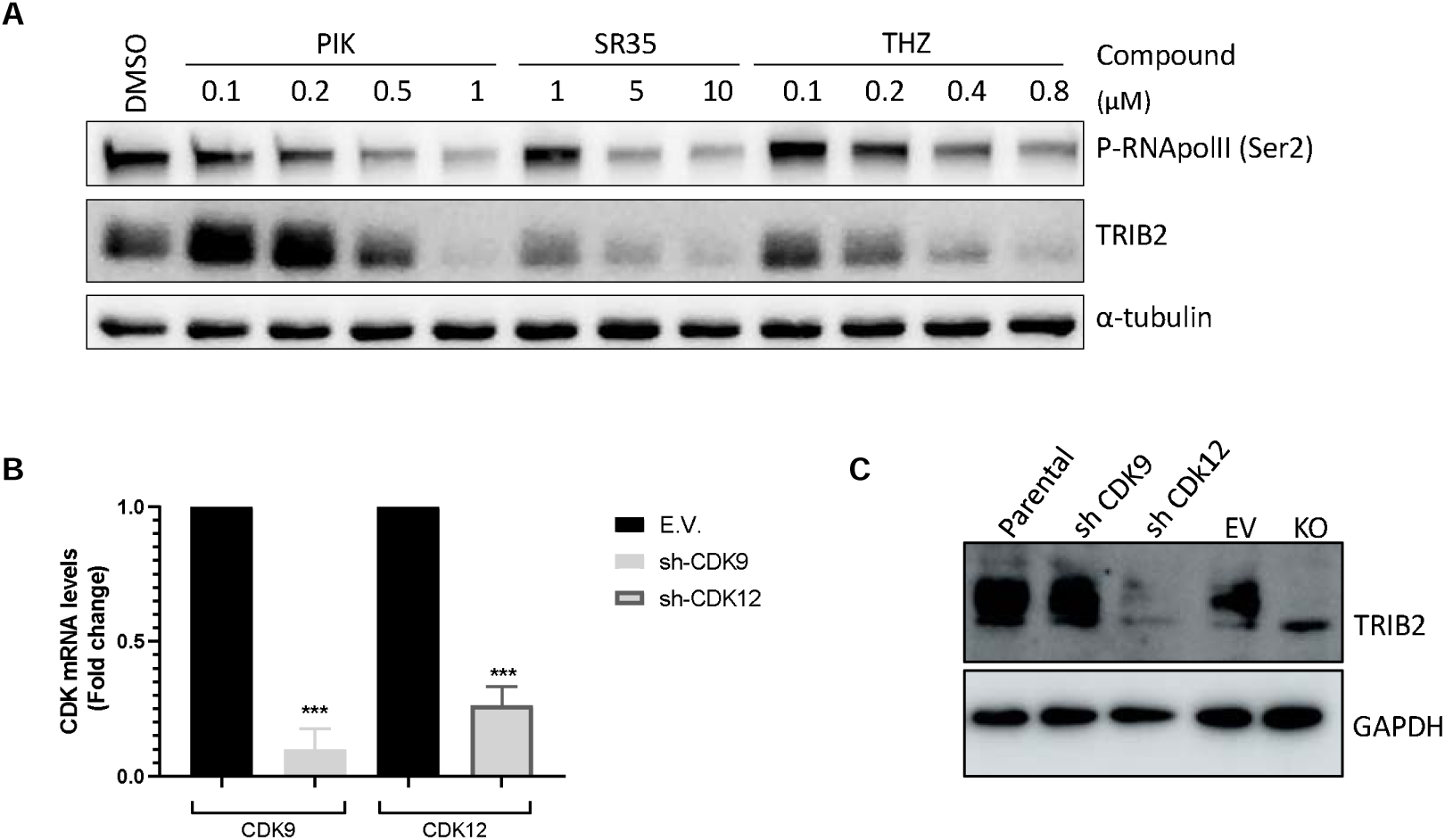
Paralleled dose-responsive reduction of Pol II CTD Ser2 phosphorylation and TRIB2 protein amounts. A) UACC-62 cells were treated with 100nM, 200nM and 500nM PIK-75, 1 µM, 5 µM and 10 µM SR-4835 and 100nM, 200nM and 500nM THZ531 for 4 hours. *In all panels, TRIB2 levels were monitored by western blot using a specific anti-TRIB2 antibody. DMSO served as the vehicle control, and* α*-tubulin was used as the loading control. B) PCR quantification of shRNA silencing of CDK9 and CDK12…….C)* These findings indicate that CDK12/13 inhibition reduces TRIB2 protein levels and that PIK-75 is a potent inhibitor of these kinases. However, because Ser2 of the RNA polymerase II CTD is also phosphorylated by CDK9, we cannot exclude the possibility that PIK-75’s effect is mediated, at least in part, through CDK9.

### PIK-75 reduces phosphorylation of the preferred CDK12/13 substrate

We next hypothesized that if PIK-75 destabilizes TRIB2 via CDK12/13 inhibition, it should also reduce phosphorylation of their preferred substrate, Ser2 of the RNA polymerase II CTD ^26^. Furthermore, the concentrations of PIK-75 and other CDK12/13 inhibitors required to reduce Pol II CTD Ser2 phosphorylation should correspond to the concentrations that trigger TRIB2 destabilization. To evaluate this, we conducted dose– response experiments in which UACC-62 cells were treated for four hours with increasing concentrations of PIK-75, or with the CDK12/13 inhibitors SR-4835 and THZ531.

As shown in Figure 7A, all three compounds, PIK-75, SR35, and THZ reduced Pol II CTD Ser2 phosphorylation in a dose-dependent manner. This reduction closely paralleled the decrease in TRIB2 protein levels. Specifically, PIK-75 reduced both TRIB2 and Ser2 phosphorylation at 0.5 and 1 µM. Similarly, SR-4835 and THZ531 showed comparable effects at 5/10 µM and 0.2/0.4/0.8 µM, respectively.

### RNAi-mediated silencing of CDK12 but not CDK9 reduces TRIB2 levels

To further investigate the underlying mechanism, we silenced CDK9 or CDK12 using shRNA and assessed the impact on TRIB2 protein levels. UACC-62 cells were transfected with shRNA constructs targeting either CDK9 or CDK12, and stable populations were generated by puromycin selection. To confirm the efficacy of the knockdowns, we performed quantitative PCR, which demonstrated a robust reduction in the expression of CDK9 and CDK12, respectively (Figure 7B). While CDK9 mRNA levels were reduced by approximately 90%, CDK9 protein levels were reduced by approximately 75%. Importantly, targeting CDK12 led to a significant reduction in TRIB2 protein levels, whereas CDK9 silencing had no effect. These results indicate that CDK12 maintains TRIB2 expression through a stabilizing mechanism, and that this function is inhibited by PIK-75.

## Discussion

TRIB2 has emerged as a promising therapeutic target in various tumor types, particularly in melanoma, to overcome therapy resistance ^3,17,27^. In this study, we present the identification of pharmacological strategies to modulate TRIB2 expression at both the transcriptional and protein levels, shedding light on the underlying molecular mechanisms. We identified four distinct pharmacological effects on TRIB2, each mediated by different molecular mechanisms. First, inhibition of the PI3K/AKT signaling pathway increased TRIB2 protein levels. Second, inhibition of the MAPK pathway reduced TRIB2 expression at the transcriptional level. Third, PLK inhibition both downregulated TRIB2 mRNA and destabilized the TRIB2 protein. Finally, CDK12/13 inhibition destabilized TRIB2 protein independently of transcription, and this effect persisted even in the presence of PI3K inhibitors, indicating a dominant mechanism. Notably, the observation that chemical agents can upregulate TRIB2 protein levels suggests their potential to drive drug resistance mechanisms and potentially elicit a feedback loop. Compounds targeting components of the insulin pathway, such as PI3K, AKT, or mTOR, reduce phosphorylation and enzymatic activity of AKT ^28^. Paradoxically, they simultaneously increase the expression of TRIB2, known to activate AKT. This could represent a potent mechanism of acquired drug resistance following treatment with these inhibitors. Further exploration of the clinical impact of this mechanism, correlating TRIB2 expression with response to these agents, is warranted. This task will become more achievable as more of these compounds enter clinical practice and patient samples become more accessible. Validation of the relationship between TRIB2 levels and resistance to these drugs would provide a strong rationale for clinical trials investigating combination therapies incorporating TRIB-targeting agents. The stabilizing effect of PI3K/AKT signalling on TRIB2 might be mediated by P70S6K which has been reported to phosphorylate TRIB2 at Ser83 ^29^. Impaired P70S6K-mediated TRIB2 phosphorylation has been shown to promote TRIB2 stabilization.

Notably, our discovery that vemurafenib and trametinib, two targeted drugs approved for treating BRAF-mutated melanomas, downregulate TRIB2 - a significant driver of therapy resistance is highly intriguing. We hypothesize that the drug-induced downregulation of TRIB2 contributes to the clinical efficacy of these agents. Investigating the relationship between TRIB2 expression and response to vemurafenib and trametinib treatment is necessary to test this hypothesis. If clinically relevant, we predict that these inhibitors would be more effective in patients with BRAF-mutated tumors that also exhibit high TRIB2 expression, compared to those with low TRIB2 levels. In this context, TRIB2 may serve as a predictive biomarker to stratify patients most likely to benefit from these inhibitors.

Additionally, our data linking PLK inhibition to TRIB2 downregulation establishes a connection between these kinases involved in eukaryotic cell division and TRIB2. Two chemically similar derivatives, BI-2536 and volasertib, were found to effectively mediate TRIB2 downregulation with high potency. Notably, while both compounds did not affect TRIB1 protein levels, they effectively reduced endogenous but not ectopically expressed TRIB3 proteins.

Regarding the bimodal effect of the PI3K inhibitor PIK-75 on TRIB2, two hypotheses are considered. At higher nanomolar concentrations, PIK-75 may inhibit other kinases or enzymes that stabilize TRIB2, or it may directly bind to TRIB2 and lead to its destabilization. CETSA experiments showed that PIK-75 did not directly bind to TRIB2. Conversely, using a combination of proteomics, chemical biology, and genetic manipulation, we identified CDK12/13 as the kinases responsible for mediating the effect of PIK-75 on TRIB2 and overriding its stabilizing effect. Notably, CDK12/13 inhibition did not alter TRIB1 or TRIB3 levels, an observation that is particularly intriguing. Together with the finding that this regulation occurs within the first 64 amino acids of TRIB2, this provides a unique opportunity to localize CDK12/13 consensus sites absent from the other Tribbles isoforms. Specifically, TRIB2 contains a stretch of serine and threonine residues, each followed by a proline, between amino acids 40 and 52, which could represent potential targets for CDK12/13-mediated phosphorylation that stabilizes the protein. While an indirect effect cannot be ruled out, the observation that treatment with these inhibitors collapses the double band of TRIB2 in western blots suggests the loss of a post-translational modification upon CDK12/13 inhibition. CDK12/13 are serine/threonine protein kinases that regulate transcriptional and post-transcriptional processes and have emerged as therapeutic targets for several types of cancer^30–32^

Our data provide additional rationale for using TRIB2 levels as a biomarker to identify patients who may benefit from CDK12/13 inhibitors.

In summary, our study highlights TRIB2 as an attractive therapeutic target in melanoma and potentially other tumor types. We have identified pharmacological approaches to manipulate TRIB2 expression at both the transcriptional and protein levels, uncovering various independent, molecular mechanisms involved. The findings suggest the potential of these compounds to influence drug resistance mechanisms and indicate the need for further investigation regarding the impact of TRIB2 on treatment response. The stabilizing effect of PI3K/AKT signaling on TRIB2 and the link between MAPK, PLK and CDK12/13 inhibition and TRIB2 downregulation provide valuable insights into TRIB2 regulation and its potential as a therapeutic target.

## Resource availability

### Lead contact

Further information and requests for resources and reagents should be directed to the lead contact, Wolfgang Link.

### Materials availability

All unique/stable reagents generated in this study are available from the lead contact with a completed materials transfer agreement.

### Data and code availability

## Author contributions

## Acknowledgements

This work was supported by a grant from Spanish Ministerio de Ciencia e Innovación/Agencia Estatal de Investigación and the European Regional Development Fund (PID2022-136654OB-I00 financed by MCIN/AEI /10.13039/501100011033 / FEDER, UE).

## Declaration of interests

Wolfgang Link is the scientific co-founder of Refoxy Pharmaceuticals GmbH, Cologne, Germany and is required by his institution to state so in his publications. The funders had no role in the design and writing of the manuscript. The rest of the authors declare no conflict of interest.

## Methods

### Chemical compounds

Table 1 lists compounds used to analyze TRIB2 regulation. All compounds were dissolved in dimethyl sulfoxide (DMSO) as stock solutions and stored at −80°C.

**Table 1:** List of compounds used to modulate TRIB2 levels.

| Name | Target | Source |
| --- | --- | --- |
| afatinib | EGFR | Selleckchem |
| BI-2536 | PLK1,2,3 | Selleckchem |
| CHIR-98014 | GSK-3α and GSK-3β | Selleckchem |
| cyclohexamide | 60S ribosomal subunit | Sigma Aldrich |
| dactolisib | panPI3K/mTOR/ATR | Selleckchem |
| GSK-461364 | PLK1 | Selleckchem |
| Laduviglusib | GSK-3$\alpha/\beta$ | MedChemExpress |
| LY294002 | PI3K (p110α, $\beta$, $\delta$) | Selleckchem |
| MG132 | Proteasome | Selleckchem |
| palbociclib | CDK4/6 | MedChemExpress |
| PI-103 | panPI3K/mTOR | Selleckchem |
| PIK-75 | PI3K (p110α, $\delta$, $\gamma$), DNA-PK, multikinase | Selleckchem |
| rapamycin | mTORC1 | LC Laboratories |
| SCH-772984 | ERK1/2 | Sigma Aldrich |
| SR-4835 | CDK12,13 | MedChemExpress |
| SPHINX31 | SRPK1 | MedChemExpress |
| TG003 | Clk1,4 | MedChemExpress |
| THZ531 | CDK12,13 | MedChemExpress |
| trametinib | MEK | Selleckchem |
| TWS119 | GSK-3β | MedChemExpress |
| vemurafinib | BRAF | Selleckchem |
| volasertib | PLK1,2,3 | Selleckchem |
| wortmannin | PI3K, PLK1,3 | LC Laboratories |

### Cell Culture

The human melanoma cell lines A375 and UACC-62, and the osteosarcoma cell line U2OS were purchased from ATCC, maintained in DMEM supplemented with 10% FBS (Sigma) and antibiotics (Gibco). Cell cultures were maintained in a humified incubator at 37°C with 5% CO2 and passaged when confluent using trypsin/EDTA.

### Plasmids and cell transfection

Expression vectors were transiently transfected into UACC-62 parental or TRIB2 KO cells with Lipofectamine 2000 transfection reagent (Thermo Fisher Scientific) following manufacturers protocol and analyzed 48 hours after transfection. We used different plasmids preceded by a constitutive promoter and tagged with myc, the complete TRIB2 sequence, the complete TRIB3 sequence and two myc-TRIB2 deletion mutants: for the N terminal (aminoacids from 64 to 353) or the C terminal (aminoacids from 1 to 308)^33^.

### siRNA-mediated silencing

A heterogeneous validated pool of siRNA against PLK2 mRNA (esiRNA, MERCK) was transiently transfected using Lipofectamine 2000transfection reagent (Thermo Fisher Scientific) for 48 hours following the manufacturer’s protocol. The effective downregulation of PLK2 and its effect on TRIB2 levels were analyzed by Western blot.

### Quantitative Real-Time PCR

To determine the transcription levels of genes affected by PIK-75, trametinib, volasertib, SCH772984, volasertib or BI-2536 treatments, cells were treated with each drug alone, 24 hours after plating, at a cell confluency of 70-90% on 6cm plates. After 6 hours of treatment, RNA was extracted (DNA, RNA and protein purification Kit, #74098 Macherey-Nagel) and then converted to cDNA (High Capacity cDNA Reverse Transcription, Applied Biosystems). Quantitative PCR was performed on 7900HT Fast Real Time PCR (Applied Biosystems) using SYBR GREEN PCR Master Mix (Applied Biosystems). Relative quantification of gene expression was determined by the 2-ΔΔCt method^34^.

### Cycloheximide (CHX) chase assays

Protein stability was assessed by using CHX chase assays. Protein biosynthesis was blocked by the inhibition of translational elongation using 50 μg/mL CHX. Subsequently, protein lysis was carried out at the indicated time points, followed by Western Blot analysis to determine TRIB2 protein degradation over time.

### Cellular Thermal Shift Assays (CETSAs)

To investigate the direct binding of small molecule compounds to their target protein, we employed the Cellular Thermal Shift Assay (CETSA), following a previously published protocol ^35^. Briefly, cells were cultured for 24 hours, washed with PBS, and protein extraction was performed using PBS supplemented with protease and phosphatase inhibitors (1mM PMSF, 1mM benzamidine, 1mM iodoacetamide, 1mM phenanthroline, 0.1mM sodium orthovanadate). Subsequently, the extracted proteins were divided into separate tubes and treated with either DMSO, 100 µM afatinib, or 100 µM PIK-75 for 30 minutes to facilitate compound-protein binding. Each treatment group was then subjected to incremental temperature increases (37, 40, 45, 50, 55°C) for 3 min. Following incubation, the samples were centrifuged at 17,000 g for 30 minutes, and the resulting supernatant was collected. Finally, 1/5 volume of Laemmli 5X buffer containing 5% β-mercaptoethanol was added to the supernatant, followed by boiling at 95°C for 5 minutes. The prepared samples were then frozen at −20°C for subsequent analysis via immunoblotting.

### Western Blot

For the preparation of whole-cell lysate, cells were harvested and lysed in lysis buffer (20mM Tris pH 7.5, 150mM NaCl, 1% Triton X-100, 50mM NaF, 1mM EDTA, 1mM EGTA, 2.5mM sodium pyrophosphate, 1mM b-glycerophosphate, 10nM Calyculan A, and EDTA-free complete protease inhibitor cocktail (PIC) (Sigma). Sample buffer was added to 1X final, and samples were boiled at 95°C for 5 minutes. Samples were resolved on 10% SDS-PAGE gels, transferred to PVDF membranes and immunoblotted according to the antibody manufacturer’s instructions. The membranes were then probed with the following primary antibodies: TRIB2 (#13533 cell signaling), α-tubulin (T9026 sigma), β-actin (A5441 sigma), PI3K (#06-195 upstate), TRIB1 (09-126 millipore), TRIB3 (Ab75846 Abcam), myc.tag (16286-1-AP proteintech), phospho-P70S6K (#9234 cell signaling), P70S6K, Erk, phospho-Erk, phosphor-RSK (ABS1849 millipore), RSK (610225 BD Biosciences), phospho-RNApol II B1 (Ser2) (Millipore 04-1571), GAPDH, Vinculin, and CDK9. Horseradish peroxidase (HRP)-conjugated secondary antibodies were added (Santa Cruz Biotechnology) at 1:5000 dilution for 1 hour at room temperature. Visualization of signal was achieved using a FUSION Solo S (VILBER) and analyzed with Evolution-Capt software.

### Proteomics

Proteomic and phosphoproteomic analyses were performed at the Proteomics facility at the National Centre for Biotechnology (CNB), Madrid, Spain. Proteins were extracted from lysates of cells treated 30min with DMSO or 500nM PIK-75 using RIPA buffer supplemented with protease and phosphatase inhibitors, followed by precipitation using the MeOH/Chloroform protocol and quantification with the Pierce 660 nm reagent. Samples underwent reduction, cysteine alkylation, and tryptic digestion. Isobaric labeling was performed using TMT reagents, labeling 50 µg of peptides per condition (4 biological replicates per condition, TMT 4×2 design) along with two internal standards (IS). The labeled peptides were pooled and split into two aliquots: Total Proteome Analysis: A 100 µg aliquot was fractionated into four fractions using high-pH reversed-phase chromatography. Phosphoproteomic Analysis: A 400 µg aliquot was desalted (Sep-Pack C18), enriched for phosphopeptides using TiO_2_, and further purified using an Oligo R3 reverse-phase polymeric column. Mass spectrometry analysis was conducted using an Orbitrap Exploris 240 system, with nano-HPLC-MS/MS separation. Peptides were ionized and analyzed in a mass range of 375–1200 m/z, with the 20 most intense precursor ions selected for fragmentation. A 90-minute LC gradient was used for total proteome analysis, while a 120-minute gradient was used for phosphoproteomics. Data analysis was performed using Proteome Discoverer v2.5 with Mascot, Sequest HT, and MSFragger search engines. Peptide identification was validated at an FDR <1%. TMT-based quantification was normalized using total peptide abundance, and statistical significance was assessed using a t-test with Benjamini-Hochberg correction.

## Notes

### Summary of Updates

This version of the manuscript has been revised to incorporate new results establishing mechanistic insight into therapeutic destabilization of the oncoprotein TRIB2

