## Supplementary material for "Pharmacological Modulation of TRIB2 Stability in Melanoma Reveals CDK12/13 as Dominant Regulators and Potential Therapeutic Targets": Suppl. Material

### Supplementary Figures

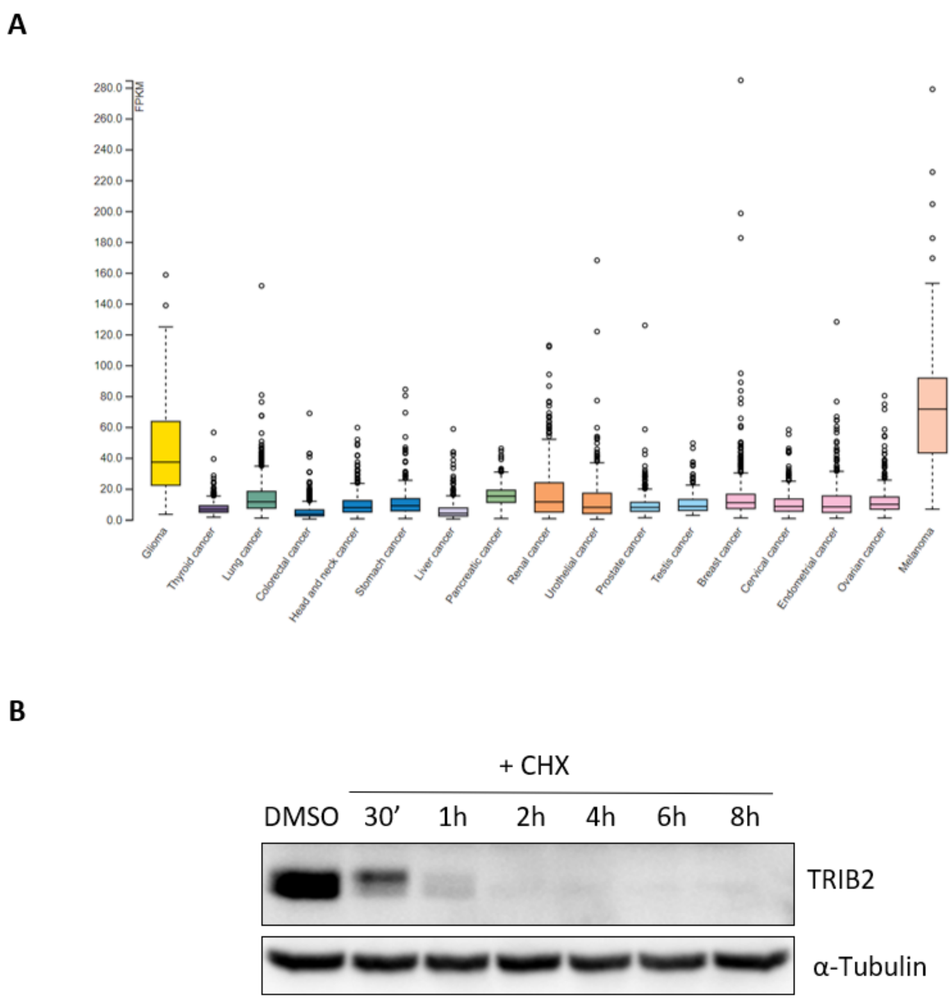


***Supplementary Figure 1*** *A) Expression of TRIB2 based on RNA-seq data from 17 cancer types from TCGA data. The TCGA RNA-seq data was mapped using the Ensembl gene id available from TCGA, and the FPKMs (number Fragments Per Kilobase of exon per Million reads) for each gene were subsequently used for quantification of expression with a detection threshold of 1 FPKM. B) Protein synthesis was blocked by Cycloheximide (CHX, 50 ug/ml) for the indicated time. The half-life of TRIB2 was measured by Western blot. α-tubulin was used as loading control.*

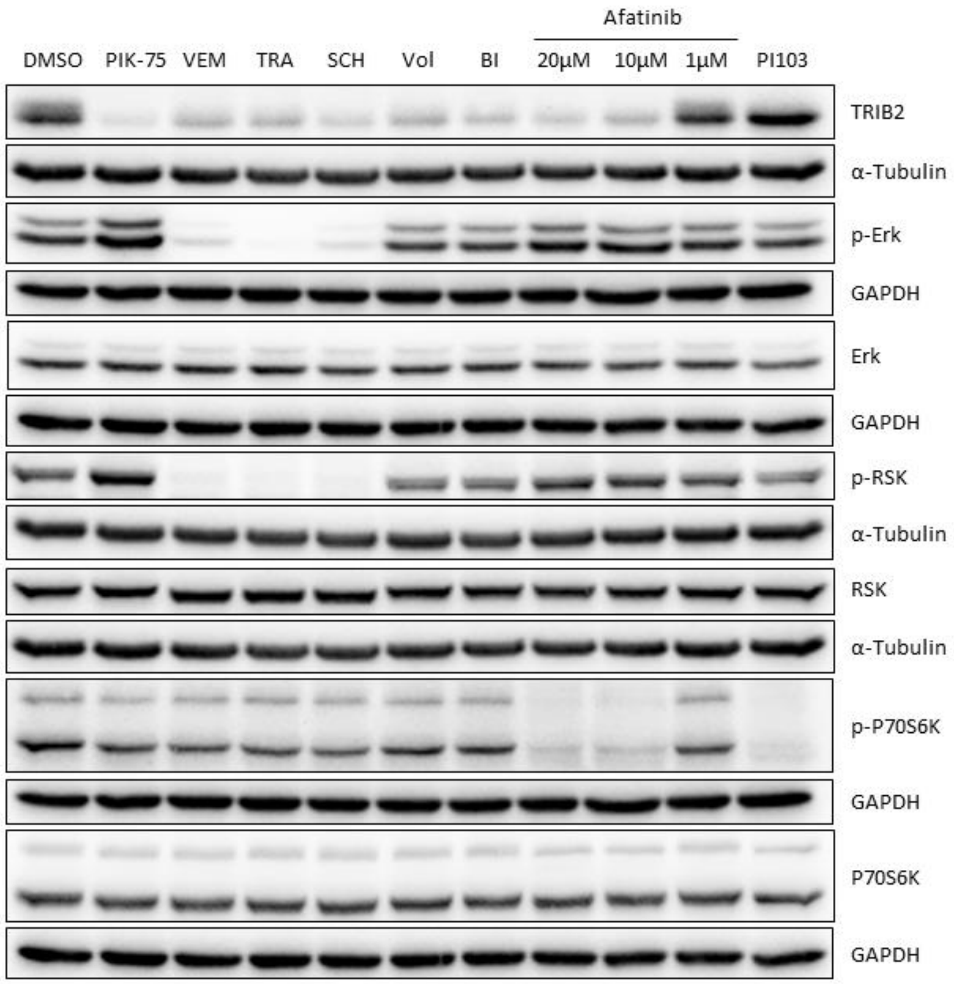


***Supplementary Figure 2*** *Effect of compounds that effect TRIB2 expression related to the MAPK pathway and PI3K pathway. p-Erk and p-RSK were used as targets of the MAPK pathway and p-P70S6K as target of the PI3K pathway. Protein levels were determined by western blot after 4h of treatment with 500nM of all compounds except afatinib which was used at the indicated concentrations. α-tubulin and GAPDH were used as loading controls.*

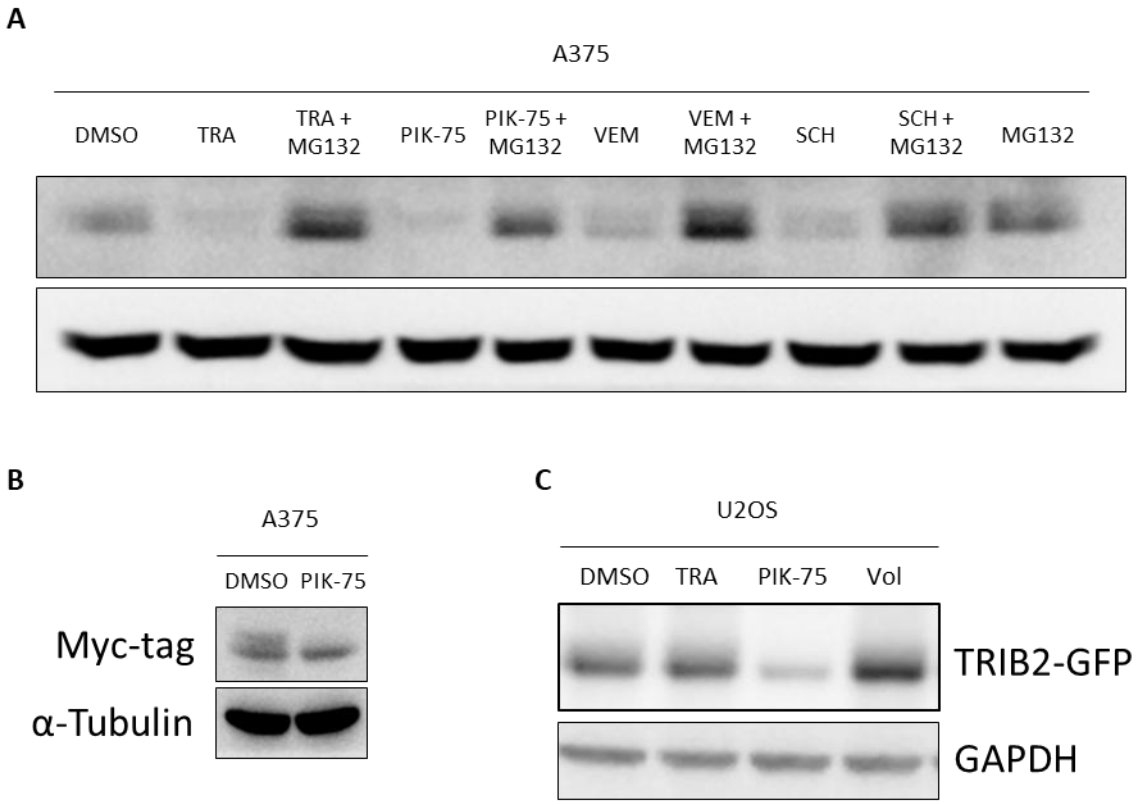


***Supplementary Figure 3*** *A) The treatment with Trametinib 500nM (TRA), PIK-75 500nM, vemurafenib 500nM (VEM), or SCH-772984 500nM (SCH) decreases the level of TRIB2 as determined by Western Blot in the A375 melanoma cell line. B) Transfected myc-TRIB2 is also downregulated by PIK-75 treatment at 500nM and 4h in the A375 cell line. C) Also, PIK-75 treatment but not Trametinib (TRA) or Volasertib (Vol) at 4h in U2OS sarcoma cell line with a constitutive expression of TRIB2 GFP reduce the TRIB2 levels. α-tubulin was used as loading control.*

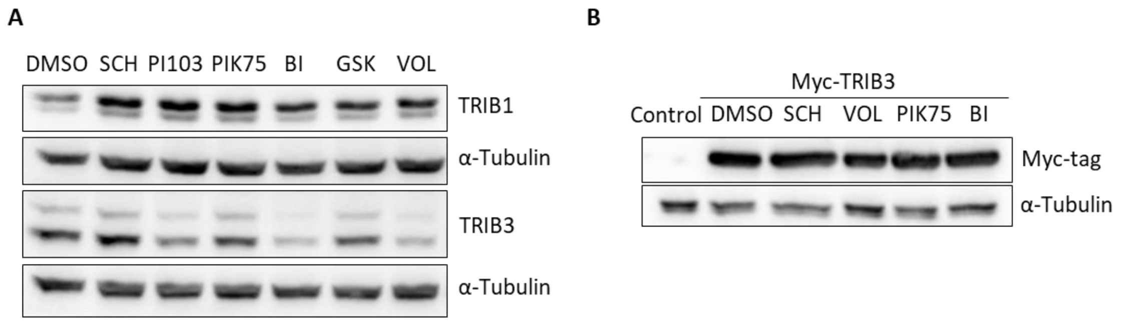


***Supplementary Figure 4* Effect of a panel of kinase inhibitors on TRIB1 and TRIB3 levels.** *A) Effect of PI-103, SCH-772984 (SCH), PIK-75, BI-2536 (BI), GSK-461364 (GSK) and volasertib (VOL) on TRIB1 and TRIB3 levels as determined by Western Blot in the UACC-62 melanoma cell line using specific antibodies against TRIB1 and TRIB3. B) Effect of SCH-772984 (SCH), PIK-75, BI-2536 (BI) and volasertib (VOL) on myc-TRIB3 ectopically expressed. Non-transfected cells were used as control and α-tubulin was used as loading control.*

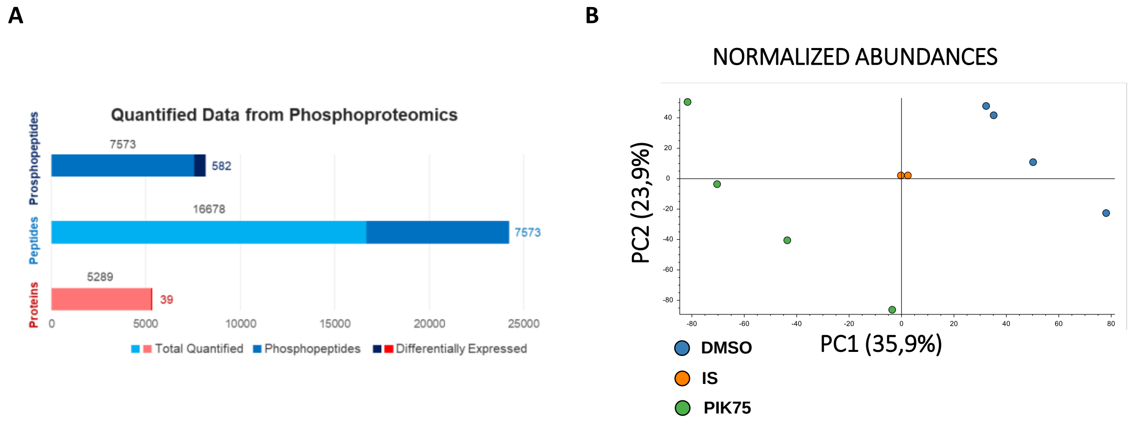


***Supplementary Figure 5* Quantified proteomics data and PCA analysis.** *A) Number of peptides and proteins identified in the proteomics and phosphoproteomics analysis of UACC62 cells treated with DMSO or PIK-75 500 nM for 30 min. First, the total proteins identified by proteomics (5,289), of which 39 show significant differences between treatments. Below is the total number of peptides identified by phosphoproteomics: 16,678, of which 7,573 are phosphopeptides. Finally, of these phosphopeptides, 583 were identified as having significant (p-value <0.05) changes >32% (log2 FC value <-0.4 or >0.4) and B) Graph of the principal component analysis (PCA) of the phosphoproteomics data.*

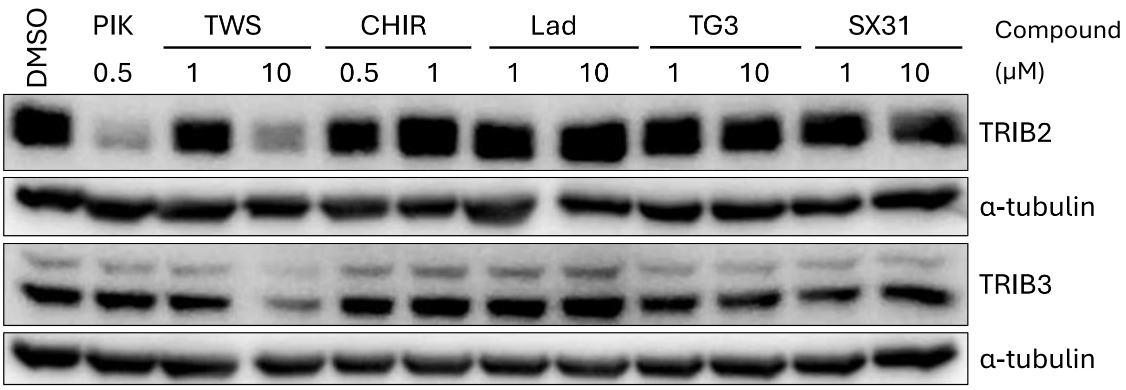


***Supplementary Figure 6*** ***Effect of specific kinase inhibitors on TRIB2 levels.*** *Effect of TWS119 (TWS), CHIR-98014 (CHIR), PIK-75, THZ531 (TH31), Laduviglusib (Lad), SPHINX31 (SX31), SR-4835 (SR35) and TG003 (TG3) on the levels of TRIB2 after 4 hours of exposure as determined by Western Blot using specific antibodies against TRIB2. Compound concentrations in µM. DMSO was used as a vehicle control and α-tubulin as loading control.*

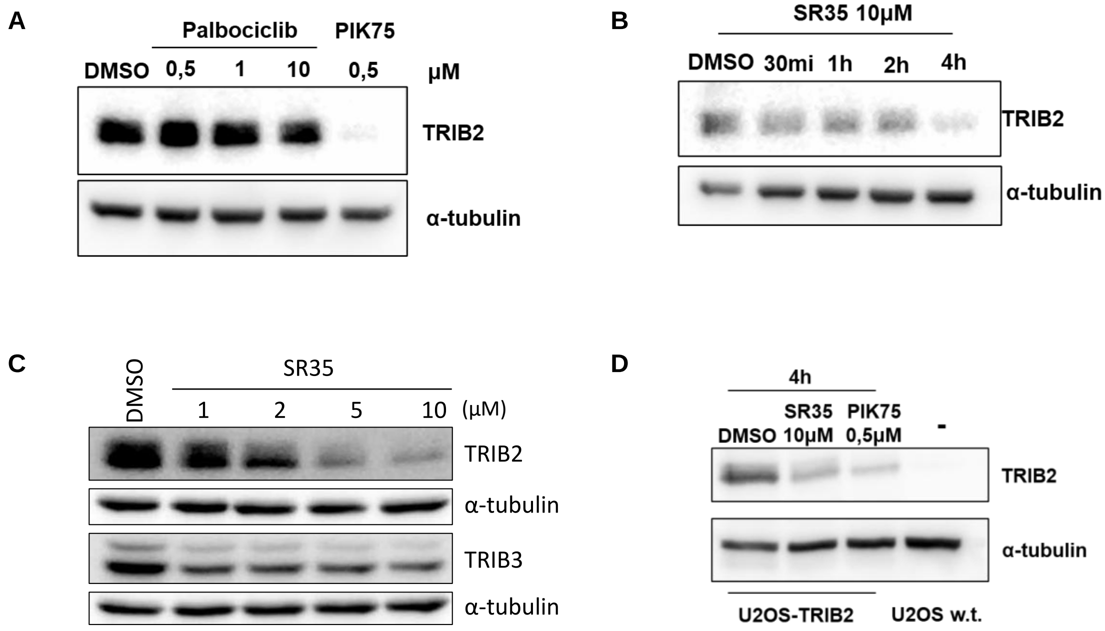


***Supplementary Figure 7. Effect of CDK inhibition on TRIB2 levels.*** *A) Effect of palbociclib on TRIB2 levels. UACC-62 cells were treated with various concentrations (µM) of palbociclib for 4 hours. PIK-75 was used as a positive control. B) Time course of TRIB2 levels following treatment of UACC-62 cells with 10 µM SR-4835 for 30 minutes (mi), 1, 2, and 4 hours (h). C) Dose-response of TRIB2 levels upon exposure to 1, 2, 5, and 10 µM SR-4835 for 4 hours. D) Effect of SR-4835 on ectopically expressed TRIB2 that cannot be transcriptionally regulated. U2OS cells lacking endogenous TRIB2 expression (U2OS w.t.) were engineered to constitutively express TRIB2-GFP (U2OS-TRIB2). U2OS-TRIB2 cells were treated with 10 µM SR-4835 for 4 hours. PIK-75 was used as a positive control. In all panels, TRIB2 levels were monitored by western blot using a specific anti-TRIB2 antibody. DMSO served as the vehicle control, and α-tubulin was used as the loading control.*
